# Recombinant surface display of lichenase on *Bacillus subtilis* spores to improve animal feed digestibility

**DOI:** 10.64898/2026.10.21.760750

**Authors:** Rohit P. James, Santosh B. Noronha

## Abstract

*Bacillus subtilis* is a well-characterised sporulating organism that has been extensively used as a probiotic in animal feed. The last few decades have seen increasing usage of feed enzymes to improve the nutrient availability of feed substrates. Lichenases are *β*-1,3-1,4-glucanases that hydrolyse mixed *β*-glucans, a major component of feed non-starch polysaccharides (NSP). Spores of *Bacillus subtilis* have been engineered to display proteins of interest on their surface. In this study, an industrially relevant lichenase, BglS was displayed on *Bacillus subtilis* spores via genetic fusion with native spore coat proteins, CotB, CotC and CotG. Among the spore-surface-displayed (SSD) constructs, CotB-BglS showed the greatest thermal stability compared to the free enzyme (*∼*30% activity remaining after 4 hours at 50°C) and retained activity after exposure to pepsin and trypsin. Moreover, CotB-BglS outperformed the free enzyme on feed substrates, achieving an 87.5% higher yield of reducing sugars from oat bran and a 150% higher yield from wheat bran. To the best of our knowledge, this is the first report of recombinant spore surface display of a lichenase. Thus, recombinant *Bacillus subtilis* spores displaying BglS can hydrolyse feed substrates and may be developed to combine feed-enzyme delivery with probiotic activity.

## 1 Introduction

Affordability and accessibility have made poultry the most widely consumed meat in the world [50]. The poultry industry runs on tight margins [65], prompting a focus on cost-cutting. This shifted attention toward feed, which remains the largest cost component of poultry production [56]. Field observations and controlled feeding trials indicated that feed was underutilised by poultry birds due to inefficient digestion [12]. Further analysis revealed the presence of non-digestible polysaccharides (anti-nutritional factors; ANFs) in the feed, which require pretreatment. Over the past few decades, the poultry feed industry has seen a significant increase in the application of exogenous enzymes to address these antinutritional effects [3]. Monogastric animals, such as broiler chickens, lack the digestive enzymes needed to release nutrients, such as phosphorus and amino acids, from feed substrates like corn [59] and wheat bran [84]. A major challenge in feed digestion is the presence of non-starch polysaccharides (NSP) [11], which constitute a significant fraction of plant-derived poultry feed ingredients [3]. Soluble NSP absorb water in the gastrointestinal tract and increase the viscosity of digesta [75]. Their presence alters the intestinal environment, reducing digestion efficiency and resulting in higher feed conversion ratios (FCR; feed intake per unit of body-weight gain) [10] and wet litter [16]. Supplementation of feed processing enzymes mitigates these issues by breaking down NSP and other ANFs such as phytate. These enzymes increase the bioavailability of nutrients and the digestibility of feed [67]. Feed enzyme supplementation also has economic and environmental benefits. It reduces the cost of separately adding bioavailable nutrients, which increases feed prices and the risk of groundwater pollution from runoff [36]. Thus, feed enzymes include NSP-degrading enzymes such as pectinases and xylanases, as well as phytases that act on phytate; these enzymes are used during feed processing and are expected to aid nutrient utilisation in the gut of the animals [46]. Lichenases are an important category of NSP enzymes. They act on mixed-linkage glucans present in substrates such as wheat bran and oat bran, cleaving *β*-1,4 glycosidic bonds in glucans containing both *β*-1,3 and *β*-1,4 linkages [22]. *β*-glucans form long soluble polymeric chains which increase intestinal digesta viscosity through water absorption and gel formation [64]. While enzymes such as lichenase improve feed utilisation, they also reduce profit margins in poultry production. Subsequent research has therefore focused on minimising enzyme-use costs through dose optimisation [14], enzyme engineering [14], and multi-enzyme cocktail development [76]. However, all these strategies focus on the addition of externally produced recombinant enzymes, which require costly purification and stabilisation steps. An alternative underutilised strategy is enzyme delivery via probiotic strains [19]. Bacillus species have been gaining popularity as probiotics due to their positive immunomodulatory effects and their ability to form highly resistant endospores that survive gastrointestinal passage [15]. Among Bacilli, *Bacillus subtilis* is a Gram-positive soil bacterium widely used as an industrial production host [83]. The microbe has been employed industrially for the production of various enzymes such as proteases [31], lipases [9] and amylases [80]. Specific *Bacillus subtilis* strains have also been used as probiotics [25].

*Bacillus subtilis* secretes proteins directly into the extracellular medium and is used to produce antigens, enzymes and other products. *Bacillus subtilis* is also the most extensively studied sporulating organism [61]. It produces endospores that are metabolically inactive and can germinate into vegetative cells under favourable conditions. Spores of *Bacillus* have been shown to be resistant to extreme changes of temperature, pH, and mechanical stress [7, 30]. Reports have suggested that they are still viable after multiple years of dormancy [69]. Nutrient germinants are sensed by receptors in the inner spore membrane, initiating the germination of dormant spores [60]. Previous research has shown that native spore coat proteins can be used as fusion partners to surface display proteins of interest. In this scheme, a native protein functions as an anchor protein, which is fused to a protein of interest in frame via a linker [27]. This approach has been used to generate spore-based vaccines [73], biocatalysts [26] and heavy metal adsorption technology [33]. Various spore surface (cot) proteins, such as CotB, CotC, and CotG have been reported for this purpose [41]. Spore surface displayed (SSD) enzymes can retain catalytic activity over repeated reuse cycles, as demonstrated for spore-displayed exochitinase [68]. Previous studies have reported increased pH and thermal tolerance of SSD enzymes compared to their free counterparts [48, 74]. Spores provide a natural scaffold for enzyme immobilisation and purification. Various categories of industrially important enzymes have been reported to be surface displayed, such as phytases [48], tyrosinases [24] and laccases [34]. Despite the widespread use of *Bacillus subtilis* as a platform for the production of exogenous feed enzymes, its spores remain underexplored as delivery systems that combine enzymatic activity with probiotic functionality. In this study, we have chosen a lichenase, BglS, native to *Bacillus subtilis 168* [77]. BglS is an industrially produced *β*-1,3-1,4-glucanase. A separate *β*-glucanase preparation produced by *Bacillus subtilis* was the subject of GRAS Notice 592, to which the FDA issued a no-questions response concerning its intended use in beer and potable alcohol production [70]. That notice does not establish GRAS status for the recombinant spores studied here. BglS was surface-displayed on spores of *Bacillus subtilis* WB800N using cot partners, CotB, CotC and CotG. This dual-purpose platform is aimed at the breakdown of soluble *β*-glucans in feed pellets and as a probiotic.

## 2 Materials and methods

### 2.1 Construction of plasmids and strains

#### 2.1.1 Vectors

The plasmid used for protein expression in *E. coli* was pET43.1b with ampicillin resistance. For expression in *Bacillus subtilis*, pBS1C, a shuttle vector, was used [55] with ampicillin resistance for selection in *E. coli* and chloramphenicol resistance in *Bacillus subtilis*.

#### 2.1.2 PCR amplification

The coat protein genes *cotB*, *cotC* and *cotG* were amplified from *Bacillus subtilis* 168 genomic DNA using primer pairs RPJ1F/R, RPJ2F/R and RPJ3F/R, respectively. Each amplicon included its native promoter, and the reverse primers encoded the flexible linker GGGGS. The amplified fragments were flanked by EcoRI and XbaI restriction sites. The *bglS* gene was amplified from the same genomic DNA using primer pairs RPJ4F/R and RPJ5F/R for insertion into pET43.1b and pBS1C, respectively. The pET43.1b insert was flanked by NdeI and EcoRI sites, and the pBS1C insert by XbaI and PstI sites. Both inserts encoded a C-terminal 6-His tag for protein purification. All primers used in this study are listed in Table 1. All restriction sites in primers are underlined.

**Table 1:**
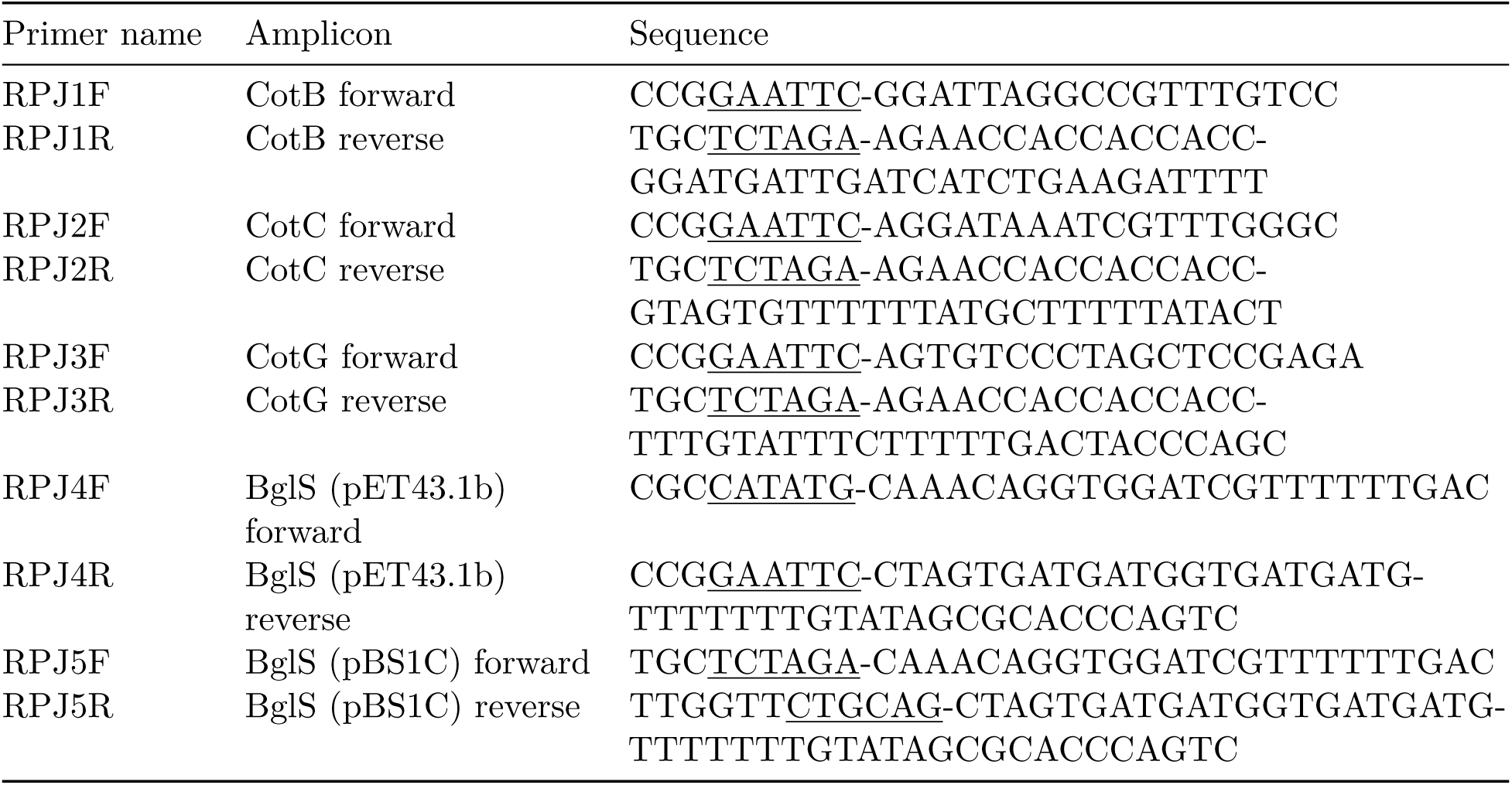
Primers used in this study.

#### 2.1.3 Cloning and transformation

After PCR, the vectors were digested with the respective restriction enzymes, and ligation reactions were set up using T4 DNA ligase. All enzymes used in cloning were procured from Thermofisher Scientific. All primers were procured from Eurofin genomics.

The ligation mix was transformed into a competent *E. coli* DH5-*α* strain. Screening was performed using colony PCR, and the selected colonies were verified by restriction digestion and sequencing of isolated plasmids. The pET43.1b construct was transformed into *E. coli* BL21 (DE3) strain. The competent cells of *E. coli* DH5-*α* and BL21 were prepared through the calcium chloride chemical competence method [58].

The screened pBS1C-based vector constructs were transformed into the WB800N strain using the natural competence protocol involving a two-step transformation as described earlier [51].

### 2.2 Preparation of Bacillus subtilis spores

Spores were prepared using Difco sporulation medium (DSM), with slight modifications to the previously described protocol [51]. Instead of the Difco broth, nutrient broth was used. Strains were inoculated into 10 mL of culture and grown overnight, then inoculated into 50 mL fresh media at 37°C and 250 rpm. Spores were harvested in 24 to 36 hours. The spores were then washed with 0.5 M NaCl and 0.5 M KCl. The washed spores were resuspended in distilled water for further use.

### 2.3 Protein purification

The BL21 cells were induced with IPTG to a final concentration of 0.2 mM and incubated for 10 hours at 25°C and 120 rpm. The cells were then harvested and lysed using a sonicator. The cell lysate was purified using Ni-NTA agarose beads from Qiagen following the manufacturer’s protocol.

### 2.4 Enzyme assay

Lichenan (Icelandic moss) obtained from Megazyme was used as the substrate. Lichenan was dissolved in 50 mM McIlvaine buffer with glycine at various pH values, and assays were performed at different temperatures to determine the optimal pH, optimal temperature and enzyme stability. Enzyme activity was calculated from the release of reducing sugars from the substrate, quantified by the DNSA method [47] using absorbance at 540 nm.

### 2.5 Estimation of enzyme loading on spores

The recombinant spores were harvested and assayed at 50°C, pH 7.0, for 30 min using 1 mg/mL lichenan as the substrate. 10^8^ spores of each construct were used for the reaction. The measured activity was then converted to enzyme mass equivalents using the specific activity of the purified BglS. This was used to estimate the amount of functional enzyme displayed on the spore surface. The calculated values were subsequently normalised and expressed as *µ*g enzyme equivalents per 10^10^ spores.

### 2.6 Protease accessibility test

The effect of proteases on SSD proteins was determined at 37°C for 1 h. Two different proteases were used in this study, pepsin and trypsin, each at 1 mg/mL. These are the commonly encountered proteases in the digestive tract of monogastric animals [48].

### 2.7 Spore recycle

Spores were assayed at 50°C and pH 7 to determine the loss of activity on reuse. After each reaction, the spores were centrifuged and resuspended in fresh substrate buffer, and the assay was repeated. Activity in the first use, shown as Control in the plot, was taken as 100%; numbered rounds indicate subsequent reuses.

### 2.8 Temperature and pH - Optimum

Spore-surface-displayed enzymes and free enzymes were compared under varying temperature (35°C– 60°C) and pH (4–8) conditions to determine their optimal values.

### 2.9 Thermal stability

Differences in thermal stability were also determined by the relative activity after incubation at 50°C. Activity was assayed every hour under optimal reaction conditions.

### 2.10 Activity on oat and wheat bran

Both oat bran and wheat bran were procured from the local market. The substrates were ground into fine powder with mortar and pestle and passed through 0.5 mm sieve. Oat bran and wheat bran were suspended separately at concentrations of 25 mg/mL and 50 mg/mL, respectively, in 50 mM McIlvaine buffer at pH 7 in 3 mL. The suspensions were assayed with DNSA before enzymatic hydrolysis and showed no detectable background sugars. Reducing sugars released during hydrolysis were quantified by the DNSA assay against a glucose standard and expressed as mM glucose equivalents. These measurements were not interpreted as the fraction of total glucan converted to glucose. One unit (U) of lichenase activity was defined as the amount of enzyme required to release 1 *µ*mol of reducing sugars per minute from lichenan under optimal assay conditions. For the assay, 1 U of free enzyme and spore-displayed enzyme equivalent to 1 U activity were used. The reactions were performed at 50°C and 250 rpm for 2 h.

### 2.11 Statistical analysis

All experiments were performed in triplicate (n = 3 independent biological replicates). Data are expressed as mean *±* standard deviation. The experimental results were plotted using OriginPro 2022b.

## 3 Results

### 3.1 Constructs and strains

The pET43.1b-bglS construct was screened using the primers RPJ4F/R, and correct construct formation was confirmed by colony PCR. Restriction digestion with NdeI and EcoRI released the expected *∼*730 bp BglS fragment (Fig. 1).

**Figure 1:**
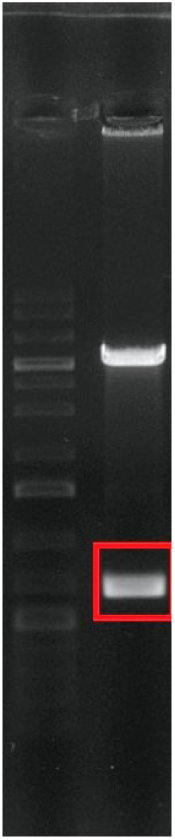
Restriction analysis of the pET43-BglS construct. Digestion with NdeI and EcoRI releases the *∼*730 bp BglS fragment.

The pBS1C-cotB-bglS construct was screened using the primers RPJ1F/5R. Colony PCR showed positive colonies with the expected band size of *∼*1870 bp. Similarly, the pBS1C-cotC-bglS and pBS1C-cotG-bglS constructs were screened using the primers RPJ2F/5R and RPJ3F/5R, respectively. The positive colonies of the former showed a band size of *∼*1144 bp and the latter *∼*1820 bp. Restriction digestion of the isolated plasmids confirmed correct assembly of all three constructs, releasing the cot-partner fragment and the cot-partner–BglS fusion fragment in the order CotB-BglS, CotC-BglS and CotG-BglS (Fig. 2).

**Figure 2:**
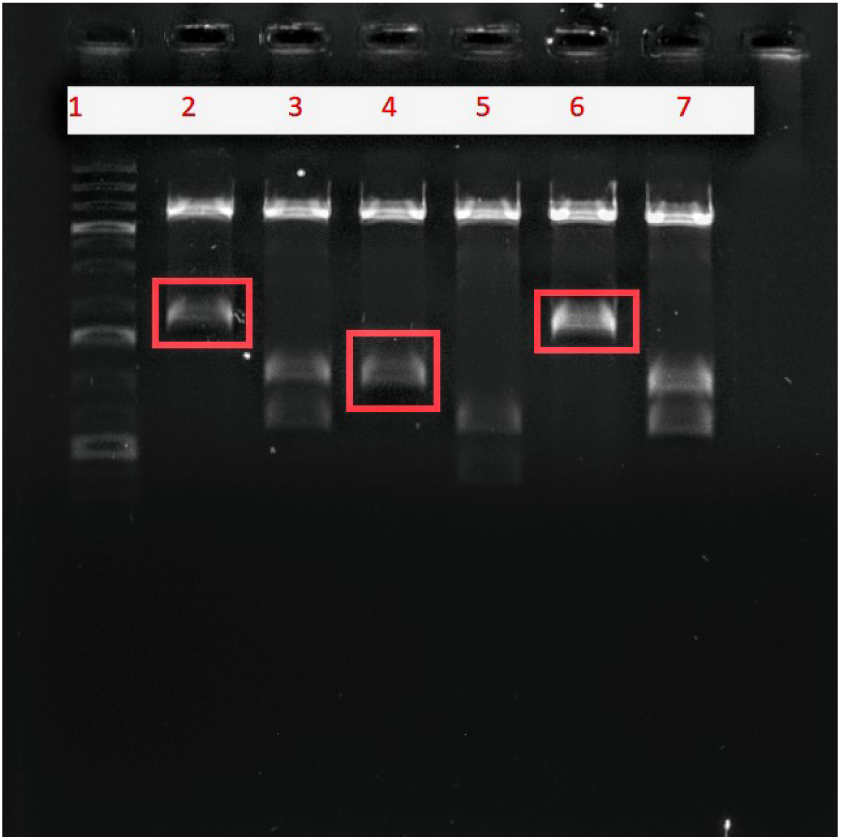
Restriction analysis of the pBS1C cot-BglS constructs. Lane 1, DNA ladder; lanes 2 and 3, CotB-BglS; lanes 4 and 5, CotC-BglS; lanes 6 and 7, CotG-BglS.

The strains used in this study are listed in Table 2.

**Table 2:** Strains used in this study.

| Strain name | Description |
| --- | --- |
| pET43-BglS (*E. coli*) | Referred to as free enzyme BglS |
| CotB-BglS (*Bacillus subtilis* sporulation strain) | Fusion between the spore coat protein CotB and BglS expressed on the spore surface. |
| CotC-BglS (*Bacillus subtilis* sporulation strain) | Fusion between the spore coat protein CotC and BglS expressed on the spore surface. |
| CotG-BglS (*Bacillus subtilis* sporulation strain) | Fusion between the spore coat protein CotG and BglS expressed on the spore surface. |

### 3.2 Protein expression and surface-display confirmation

SDS-PAGE of the pET43-BglS expression construct is shown in Fig. 3.

**Figure 3:**
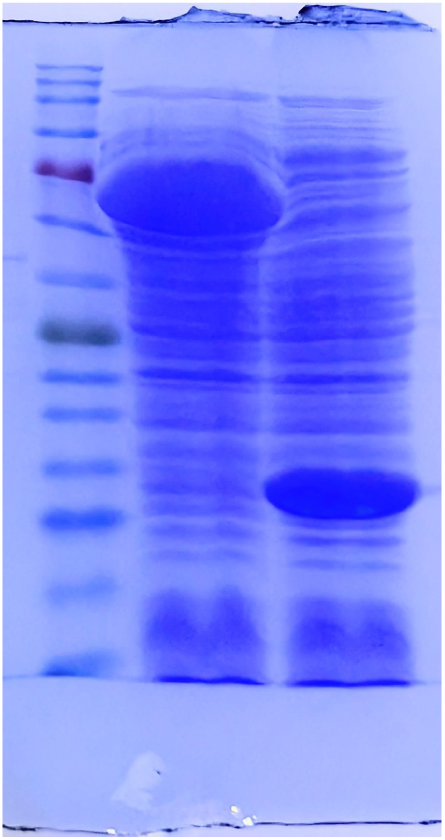
SDS-PAGE of pET43-BglS. Lanes: 1, protein ladder; 2, negative control (empty pET43-NusA tag); 3, pET43-BglS under IPTG induction.

### 3.3 Protease accessibility test

As shown in Fig. 4, incubation with the proteases pepsin and trypsin reduced the activity of all spore-displayed constructs relative to the untreated control. Trypsin caused the greater loss of activity, and the CotG-BglS construct showed the largest decline.

**Figure 4:**
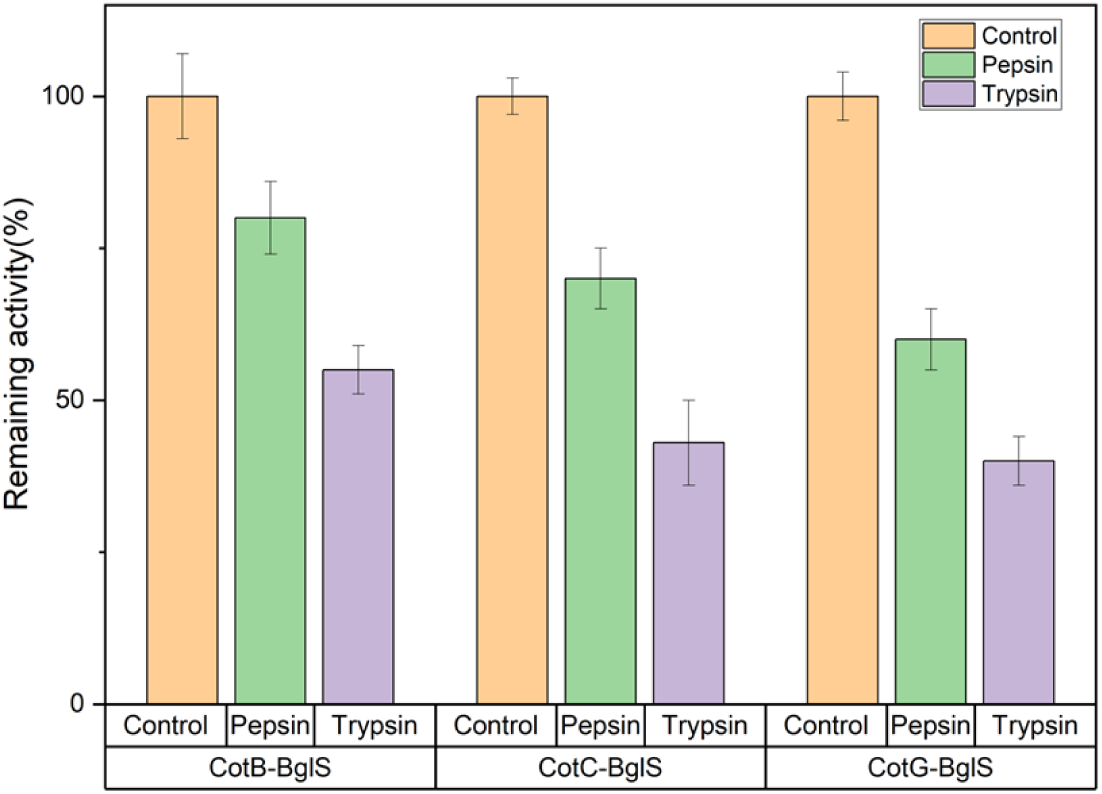
Activity of spore-displayed BglS after treatment with pepsin or trypsin for 1 h at 37°C, relative to the untreated control (100%). Quantitative plots in this paper show mean *±* SD from three independent biological replicates (n = 3).

### 3.4 Estimation of enzyme loading on spores

The recombinant spores (10^8^) were harvested and assayed at 50°C, pH 7.0, for 30 min using 1 mg/mL lichenan as the substrate.

All three fusion constructs showed activity on lichenan. The control strain, non-recombinant spores of WB800N, also exhibited some detectable background activity on lichenan. However, lichenan is a model substrate; activity may differ with other substrates. Purified BglS exhibited a measured specific activity of 2100 U/mg, consistent with previously reported values [18]. Using this value, the activity measured for each construct was back-calculated to the corresponding amount of functional enzyme displayed per 10^10^ spores, as shown in Fig. 5. CotB-BglS showed the highest loading at 3.5 *µ*g per 10^10^ spores, followed by CotG-BglS at 2.8 *µ*g per 10^10^ spores and CotC-BglS at 1.3 *µ*g per 10^10^ spores. Relative to the background activity of the control spores, the displayed enzyme density of the engineered constructs corresponded to 9-fold, 7.5-fold, and 3.3-fold higher activity for CotB-BglS, CotG-BglS, and CotC-BglS, respectively. These values represent the functional, catalytically active fraction of displayed BglS, as they are derived from measured activity.

**Figure 5:**
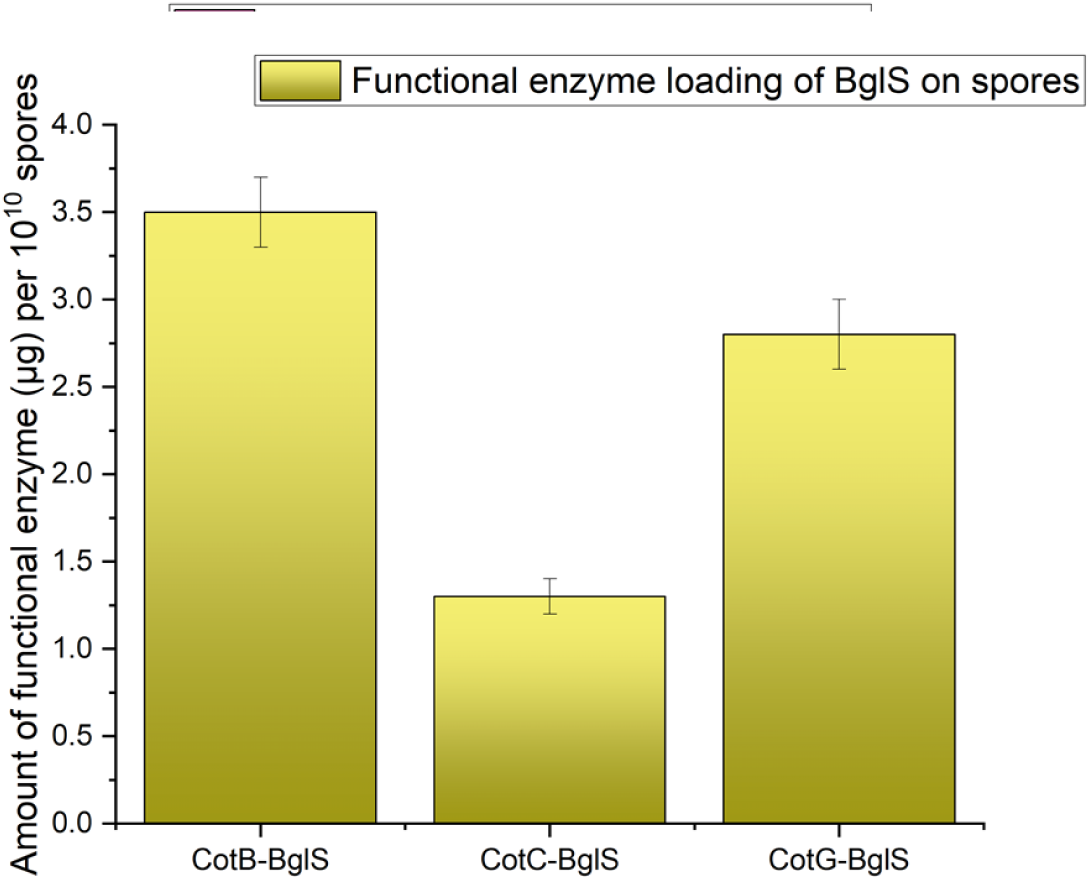
Activity-based functional loading of the three recombinant BglS constructs, expressed as *µ*g enzyme equivalents per 10^10^ spores. Activity was measured with 1 mg/mL lichenan at 50°C and pH 7 for 30 min and converted using the purified enzyme specific activity of 2100 U/mg.

### 3.5 Characterisation of free and spore displayed enzymes

#### 3.5.1 pH optimum

As illustrated in Fig. 6, the pH profiles of free and spore-displayed BglS were similar across pH 4–8, with no observable shift in the optimum after display. The free and spore displayed BglS showed a pH optimum of 7. At an acidic pH of 4, only 23–27% of peak activity remained. An increase in pH to 8 resulted in a reduction of activity to approximately 56–63% across the free enzyme and spore-displayed constructs.

**Figure 6:**
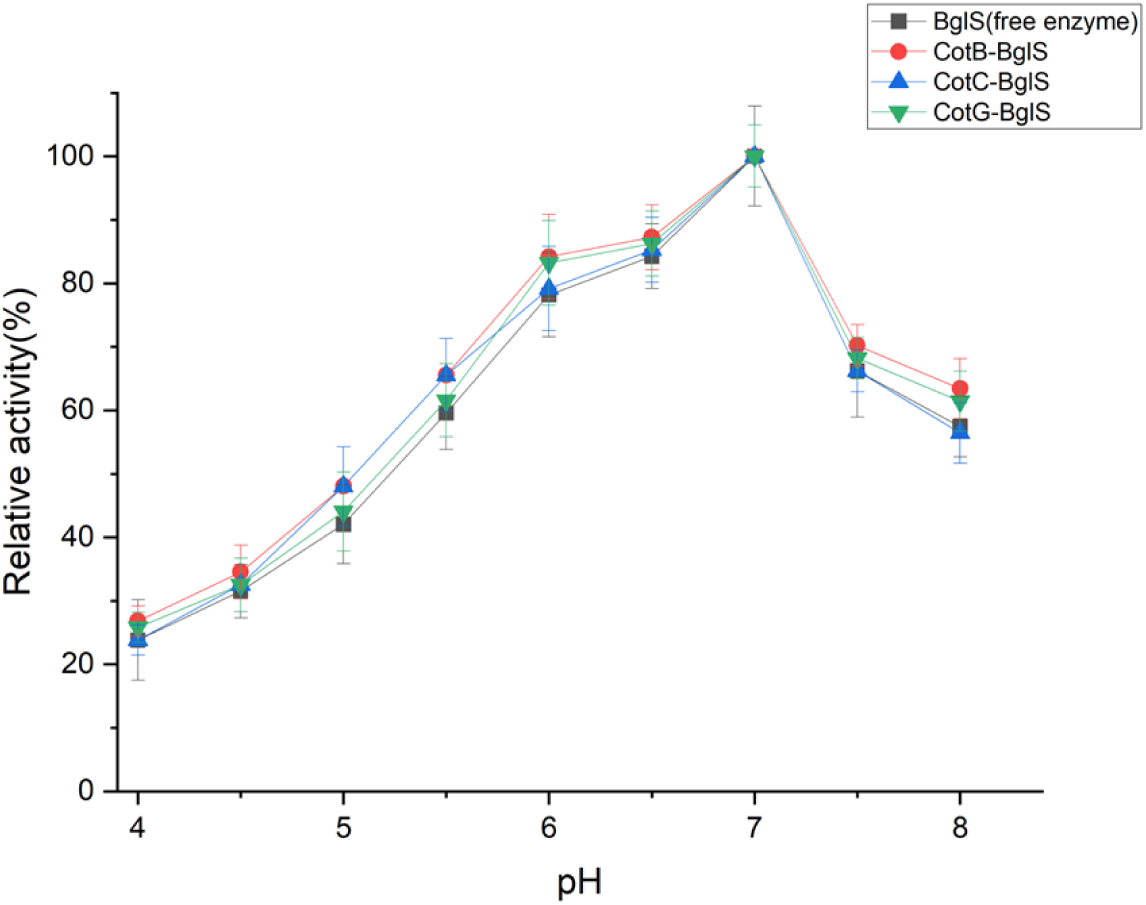
pH profiles of free BglS and the CotB-, CotC- and CotG-BglS spore constructs on lichenan over pH 4–8. Activity is expressed relative to the maximum for each preparation (100%).

#### 3.5.2 Temperature optimum

Enzyme activity was assayed over a broad temperature range (35°C–60°C). As indicated by Fig. 7, the temperature profiles of both free and spore-displayed BglS showed a temperature optimum of 50°C. While there was no shift in the optimum, CotB-BglS construct displayed enhanced activity retention at higher temperatures with *∼*76% peak activity at 60°C. All constructs exhibited similar activities at lower temperatures (*∼*77–80% activity at 35°C).

**Figure 7:**
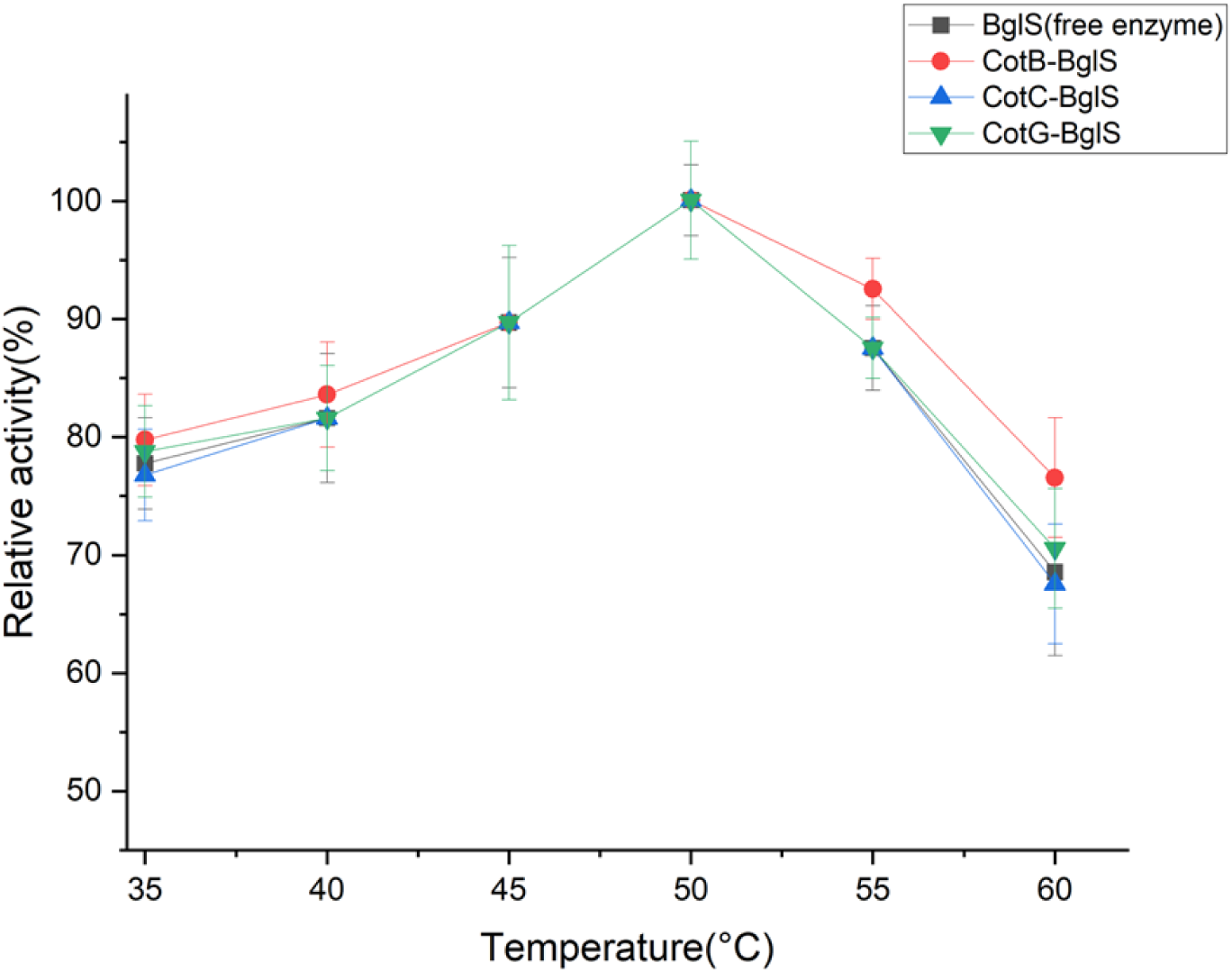
Temperature profiles of free BglS and the CotB-, CotC- and CotG-BglS spore constructs on lichenan over 35–60°C. Activity is expressed relative to the maximum for each preparation (100%).

#### 3.5.3 Thermal stability

To evaluate thermal stability, both the free and spore-displayed BglS were incubated at 50°C for 4 hours and assayed hourly. As shown in Fig. 8, CotB-BglS, CotG-BglS and CotC-BglS showed increased thermal stability compared to the free enzyme. They retained *∼*30%, *∼*24% and *∼*17% of activity, respectively, after 4 hours. In comparison, the free enzyme had no detectable activity after 4 hours.

**Figure 8:**
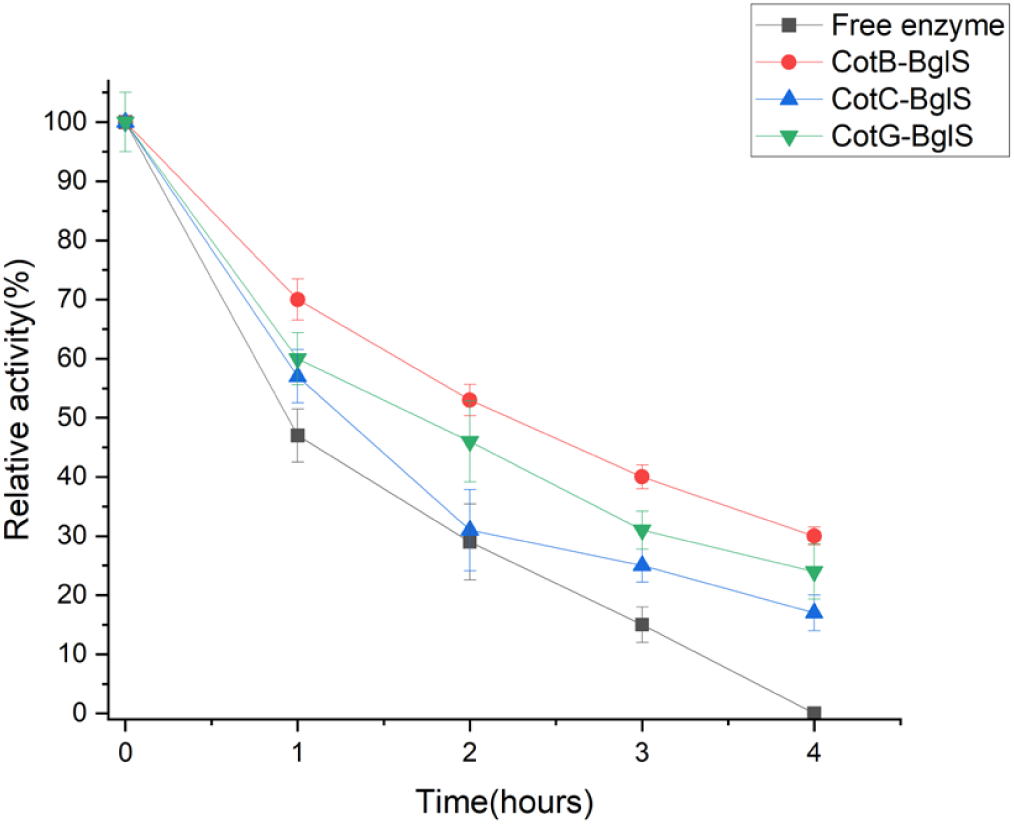
Thermal stability of free and spore-displayed BglS during incubation at 50°C for 4 h. Activity was assayed hourly and expressed relative to initial activity (100%).

### 3.6 Spore recycle

Spore-displayed BglS was tested over repeated reaction cycles under optimal assay conditions, with activity in the first use taken as 100%. As shown in Fig. 9, all three constructs demonstrated recyclability, remaining catalytically active after three rounds of reuse. CotB-BglS spores retained the maximum activity at 20%, followed by CotG-BglS (13%), and CotC-BglS showed the steepest decline with only 10% activity remaining.

**Figure 9:**
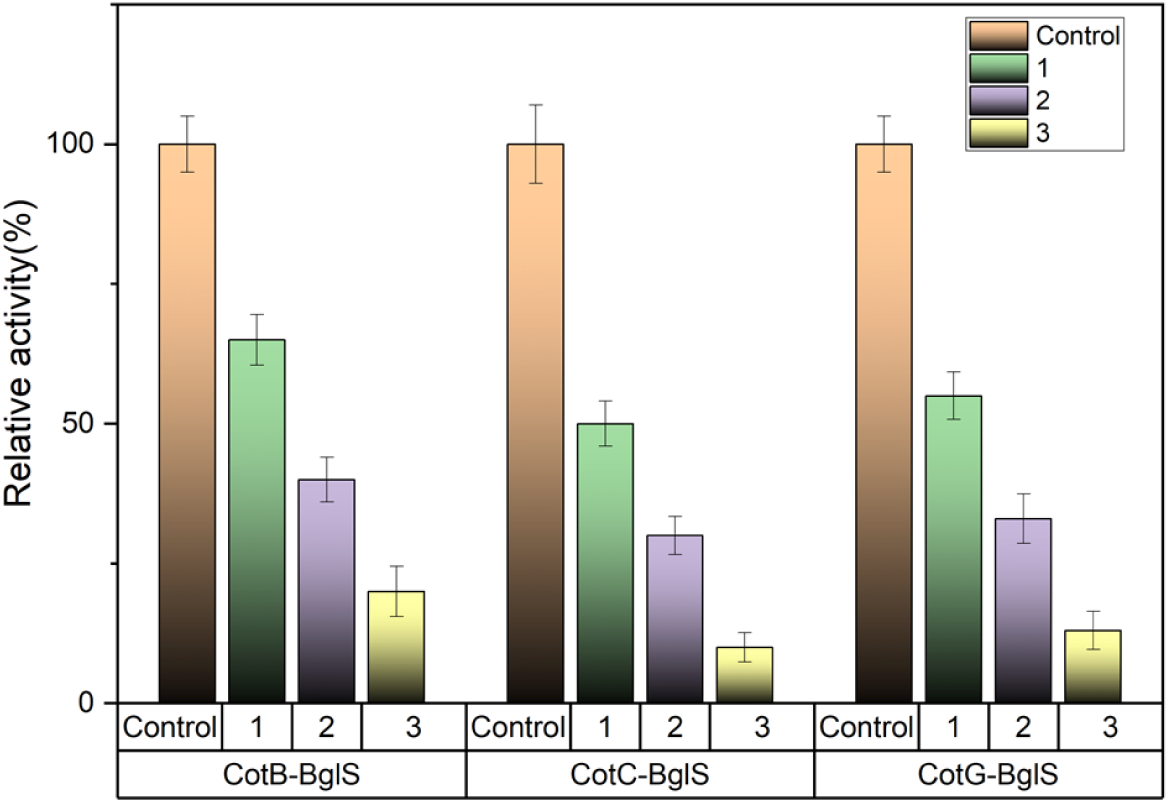
Reuse of spore-displayed BglS at 50°C and pH 7. Control is the first use (100% activity); rounds 1–3 indicate subsequent reuses, with spores recovered by centrifugation and resuspended in fresh substrate buffer between reactions.

### 3.7 Activity on feed substrates

Fig. 10 compares the activity of free and spore-displayed BglS on oat bran. The assay was performed in a 3 mL volume using the same spore-displayed enzyme equivalents (1 U) of all three spore constructs and free enzyme. CotB-BglS and CotG-BglS produced higher yields than the free enzyme. CotB-BglS gave the highest yield after a 2-hour reaction time, releasing 7.5 mM of glucose equivalents. CotG-BglS and the free enzyme released 5.8 mM and 4 mM glucose equivalents, respectively.

**Figure 10:**
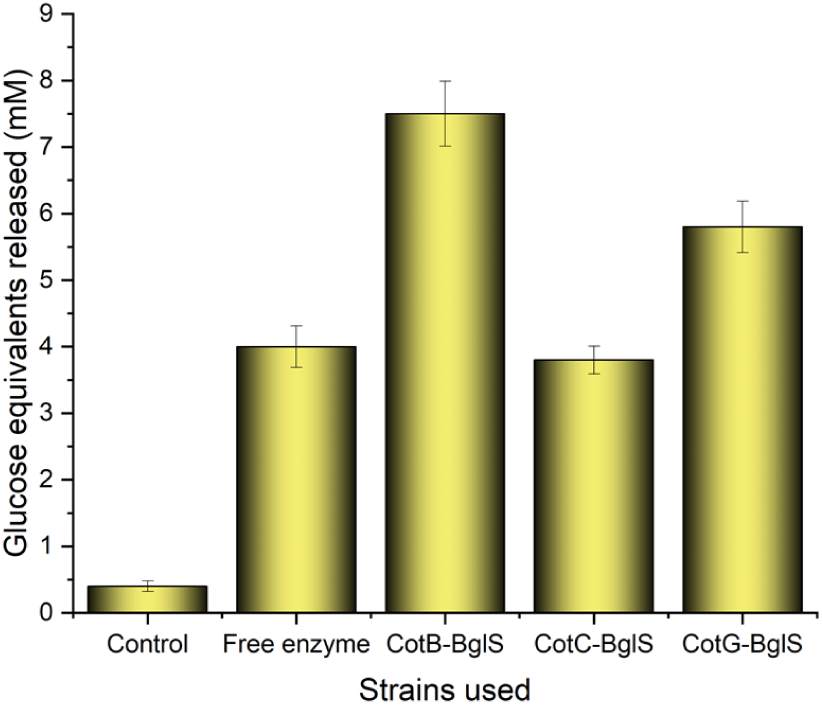
Reducing sugars released from oat bran (25 mg/mL) by free and spore-displayed BglS. Each preparation supplied 1 U activity in 3 mL of 50 mM McIlvaine buffer, pH 7, and was incubated at 50°C and 250 rpm for 2 h. DNSA measurements are expressed as mM glucose equivalents.

Similarly, Fig. 11 confirms that all four constructs are active on the substrate, wheat bran. Here again, CotB-BglS gave the highest yield, releasing 5 mM of glucose equivalents after 2 hours, followed by CotG-BglS (4 mM). However, unlike oat bran, the CotC-BglS (2.5 mM) on wheat bran performed better than the free enzyme (2 mM).

**Figure 11:**
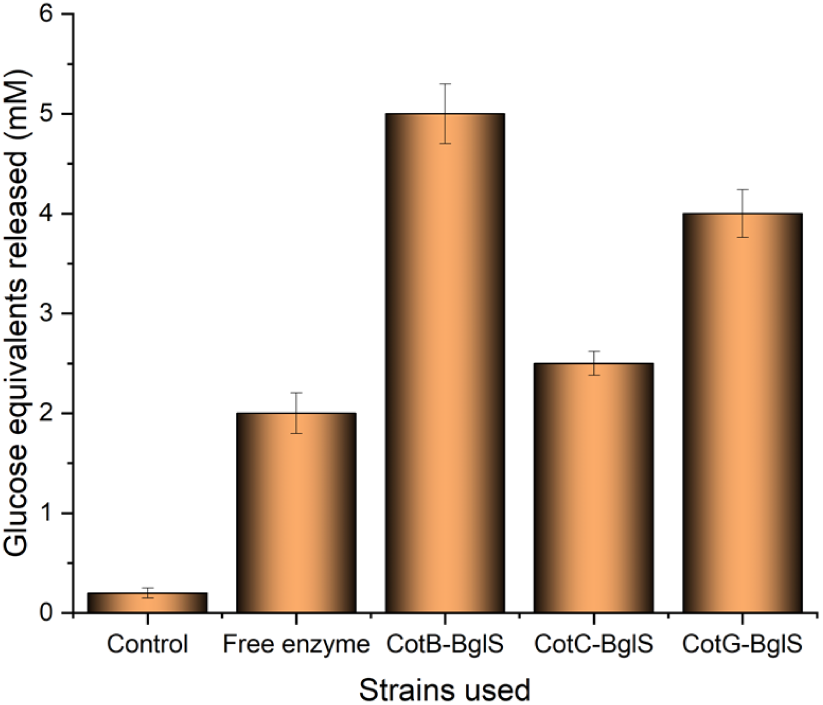
Reducing sugars released from wheat bran (50 mg/mL) by free and spore-displayed BglS. Each preparation supplied 1 U activity in 3 mL of 50 mM McIlvaine buffer, pH 7, and was incubated at 50°C and 250 rpm for 2 h. DNSA measurements are expressed as mM glucose equivalents.

## 4 Discussion

The valuation of the animal feed additives market is projected to reach US $56 billion by 2027 [63]. Feed enzymes break down NSP and ANFs. Incorporating feed enzymes into the diet improves feed conversion and reduces digestive problems, especially in monogastric animals such as chickens. ANFs such as phytic acid in feed are broken down by phytases, increasing feed nutrient bioavailability. This combined effect enhances poultry health and meat quality [5]. However, the poultry feed industry is price-sensitive, where the added costs of enzyme production, storage, and integration can be prohibitive. This study examines recombinant *Bacillus subtilis* WB800N spores as a platform for feed enzyme delivery. Probiotic supplementation improves poultry’s immune profile, reducing reliance on antibiotics [81]. Specific *Bacillus subtilis* strains are used as probiotics in the poultry industry [45]. Combining lichenase activity with a suitable probiotic strain could potentially improve feed digestibility and poultry health. BglS is a native lichenase secreted by *Bacillus subtilis* 168. Lichenases have significant industrial potential but also constraints, such as thermal stability issues, that hinder their widespread adoption [6].

In this study, BglS was surface-displayed using the recombinant approach. Fusion constructs were made with cot-partners, CotB, CotC and CotG and transformed into a protease-deficient *Bacillus subtilis* WB800N strain. Lichenase activity was detected in all three spore constructs. The reduction in activity of all constructs following protease treatment indicates that BglS is localised on the spore surface, since proteases cannot access the spore interior. The greater susceptibility of CotG-BglS may reflect greater accessibility to proteases. The activity-based loading estimates, which were severalfold above the background of non-recombinant spores, confirm that the displayed BglS is the principal contributor to the measured activity.

BglS was displayed and characterised across all three of the most commonly used outer-coat anchors (CotB, CotC and CotG). Most spore-display studies report a single anchor; the closest comparison displayed a keratinase on two anchors and likewise found CotG effective [74]. This three-anchor characterisation allows the anchor-dependent differences in activity, thermal stability and recyclability to be observed directly.

Immobilised enzymes often exhibit altered catalytic properties, such as a shift in optimal pH and temperature, due to changes in the local microenvironment surrounding the enzyme, including electrostatic interactions and diffusion [43]. Similar effects have been reported previously in *Bacillus subtilis* SSD systems, where anchoring enzymes to the spore coat may influence their behaviour [27].

In this study, SSD constructs showed no appreciable differences in activity over the pH range assayed, with an observed pH optimum of 7, similar to that of the free enzyme BglS. Previous studies comparing SSD enzymes with their free counterparts have reported increases [78], decreases [21], and retention of the pH optimum [48], consistent with observations reported here. Similarly, the temperature optimum also remained unchanged at 50°C for all constructs. However, CotB-BglS fusion displayed slightly higher activity (*∼*76% at 60°C) at elevated temperatures compared to the free enzyme (*∼*68%). Previous studies on SSD enzymes have noted increases [40], decreases [8] and retention of temperature optimum [72] as observed in this study.

Previous SSD studies have suggested that the spore coat provides a natural immobilisation scaffold in which anchored enzymes exhibit reduced conformational flexibility, thereby conferring a protective effect against thermally induced unfolding [8, 53, 57]. All SSD constructs showed improved thermal stability compared to the free enzyme. After 4 hours, CotB-BglS retained *∼*30% activity, followed by CotG-BglS (*∼*24%) and CotC-BglS (*∼*17%), whereas the free enzyme had no detectable activity. The retention of activity under prolonged thermal incubation is consistent with several reports indicating improvement of thermal stability due to spore surface-display [40, 68, 74, 78]. While lichenases are gaining attention as feed additives, laundry detergent additives, and for biofuel production, many such enzymes are unsuitable for industrial use due to their lack of thermal stability [79]. Previous attempts to improve the thermal stability of the lichenase BglS used various immobilisation methods such as SpyCatcher [71] and ELP-mediated silicification [42]. The current study demonstrates improved thermal stability at 50°C compared to the free enzyme. This supports further assessment for feed processing, although the stability measured at 50°C does not establish survival under the higher temperatures encountered during feed pelleting [20]. However, further stability improvements are required for industrial operations at higher temperatures, such as lignocellulose breakdown.

The CotB-BglS construct exhibited the highest activity per spore and the best recyclability among the three constructs, retaining 20% activity after three rounds of recycling, followed by CotG-BglS (13%). Conversely, CotC-BglS demonstrated the lowest recyclability, with only 10% activity remaining after the third recycle. Overall, CotC-BglS showed the lowest activity, thermal stability, and recyclability among the three constructs, though it exhibited higher activity than the free enzyme on wheat bran.

The spore-displayed constructs demonstrated activity on feed substrates, specifically oat and wheat bran. Both substrates contain soluble *β*-1,3-1,4-glucans, whose long polymeric chains increase the viscosity of digesta in the gastrointestinal tract. By depolymerising these glucans, BglS could reduce digesta viscosity and improve access to existing nutrients rather than increase the nutrient content of the feed. Effects on nutrient absorption and gut transit remain to be established for this system. CotB-BglS achieved maximum yield from oat bran (7.5 mM) and wheat bran (5 mM). CotG-BglS followed, producing 5.8 mM on oat bran and 4 mM on wheat bran. Notably, CotC-BglS outperformed the free enzyme on wheat bran (2.5 mM) but had a lower yield on oat bran (3.8 mM). Overall, enzymatic activity was greater for all constructs on oat bran than on wheat bran. These results suggest that immobilisation on the spore surface may improve enzyme stability compared to the free enzyme. These findings support further testing of spore-displayed BglS for feed processing. These results are based on in vitro assays with isolated bran substrates and have not yet been validated in formulated feed or in vivo.

Both substrates are rich in amino acids [2, 29]. The presence of amino acids, such as alanine, can trigger the germination of spores [52]. Vegetative cells were observed via microscopy of the oat and wheat bran reaction mixtures, indicating spore germination. Increasing incubation time beyond 2 hours achieved no appreciable yield increase (data not shown). Germination and BglS inactivation may have contributed to activity loss, but their contributions were not determined. The observed germination does not establish probiotic functionality. The probiotic potential discussed here is based on previous reports on *Bacillus subtilis* and was not assessed for these constructs. Probiotics are beneficial when therapeutic minimum colony-forming units (CFUs) are delivered to the colon [62]. For probiotic strains, ingestion as spores offers distinct advantages over vegetative cells. The highly resistant endospores ensure survival under adverse conditions, such as the acidic pH of the stomach [4]. Live probiotics generally overcome this limitation by high dosage [62] or the selection of acid-resistant strains [35]. Several studies report the usage of *Bacillus subtilis* endospores as probiotics, demonstrating increased stability [44], enhanced shelf life [17], and beneficial immune modulation [38]. *Bacillus subtilis* spores have also been demonstrated to undergo germination in the gut [13] with detection of spores in the gastrointestinal tract up to five days post ingestion [37]. Furthermore, it has been demonstrated that endospore-based probiotics undergo repeated cycles of germination and sporulation in the gut [66]. The activity demonstrated here on isolated bran substrates supports the potential of lichenase-displaying *Bacillus subtilis* spores for feed processing. Such spores could potentially be used alongside probiotic supplementation, but their compatibility with existing processing and storage conditions has not been assessed. Spore-displayed lichenase might reduce the need for separately supplied lichenase if sufficient activity is retained under feeding conditions. Any reduction in spore dosage would also depend on persistence and enzyme activity in the gut, neither of which was measured for these constructs. These possibilities require validation in feed matrices and in vivo.

To the best of our knowledge, this is the first report of recombinant spore surface display of a lichenase. CotB, CotC and CotG are the most frequently reported anchors [82]. While CotB was found to be the most suitable anchor for BglS fusion, functional expression was achieved with all three coat anchors, including CotC and CotG fusions. Therefore, this study identified BglS as a promising new reporter candidate, exhibiting activity with all three major cot partners. Established spore-display reporters such as eGFP report protein presence through fluorescence [32], and *β*-galactosidase has been used for native enzyme display [53]. BglS complements these by providing a direct, activity-based readout: as a natively encoded enzyme assayed by a simple, inexpensive DNSA method, it allows functional display to be quantified and compared across cot partners without specialised detection. The ready availability of the native gene template further facilitates rapid cloning and expression. BglS can therefore be used to validate spore surface display constructs and to assess the suitability of new anchor proteins as fusion partners through activity-based quantification. Despite their potential, regulatory oversight of GMOs remains a major challenge for recombinant spore surface-displayed enzymes. The persistence of viable recombinant spores could raise concerns about public acceptance, potential environmental and food hazards [49]. Consequently, recent research has focused on the development of bio-containment strategies, such as thymineless death (thymine auxotrophy), to prevent microbial escape [23].

## 5 Conclusion

The CotB-BglS construct exhibited enhanced thermal stability, retaining *∼*30% residual activity after 4 h at 50°C, a desirable property for animal feed processing applications. Spore-displayed BglS retained activity after protease treatment, consistent with a possible protective contribution of the spore coat. Resistance to common gastrointestinal proteases such as trypsin and pepsin suggests that the spore-displayed enzyme may retain activity under gastrointestinal conditions [48]. *Bacillus subtilis* has already been reported to undergo multiple rounds of sporulation, germination and persistence in the gut of animals such as broilers and pigs [37, 39]. *Bacillus subtilis* is a known probiotic that improves the meat quality [1] and immunity [54] of broiler chickens. CotB-BglS also showed higher yields on actual feed substrates such as oat bran and wheat bran, producing 87.5% higher reducing sugar yield from oat bran and 150% higher yield from wheat bran compared to the free enzyme. Therefore, this study demonstrates that *Bacillus subtilis* spores can be augmented with lichenase activity for in vitro hydrolysis of feed substrates. Improved animal feed digestibility and probiotic functionality remain potential applications rather than demonstrated outcomes of this study. Probiotic potential in *Bacillus subtilis* is known to be strain-dependent, depending on variations in germination efficiency, metabolite production and host interactions. Therefore, further studies using established probiotic strains such as C-3102 [28], evaluating *in vivo* feed digestibility and probiotic efficacy are necessary to establish the potential of this system for animal feed applications.

## Acknowledgements

The authors thank the current and former members of the Protein Engineering Lab for helpful discussions and technical assistance, and the Department of Chemical Engineering, Indian Institute of Technology Bombay, for providing the research facilities used in this work.

## Statements and Declarations

### Author contributions

R.P.J. and S.B.N. conceived and designed the study. R.P.J. performed the experiments, analysed the data, and wrote the manuscript. S.B.N. supervised the project, acquired funding, and reviewed and edited the manuscript. Both authors read and approved the final manuscript.

### Ethics approval

Not applicable. This study did not involve human participants or animals.

### Consent to participate / Consent for publication

Not applicable.

### Data availability

The data supporting the findings of this study are available from the corresponding author upon reasonable request.

